# Allosteric deregulation of phenylalanine biosynthesis evolved with the emergence of vascular plants

**DOI:** 10.1101/834747

**Authors:** Jorge El-Azaz, Francisco M. Cánovas, Belén Barcelona, Concepción Ávila, Fernando de la Torre

**Author notes:** **Corresponding authors**: Francisco M. Cánovas, Fernando de la Torre.

## Abstract

Phenylalanine (Phe) is the precursor of essential secondary products in plants. Here we show that a key, rate-limiting step in Phe biosynthesis, which is catalyzed by arogenate dehydratase (ADT), experienced allosteric de-regulation during evolution. Enzymes from microorganisms and type-I ADTs from plants are strongly feedback-inhibited by Phe, while type-II isoforms remain active at high levels of Phe. We have found that type-II ADTs are widespread across seed plants and their overproduction resulted in a dramatic accumulation of Phe in planta, up to 40-times higher than those observed following the expression of type-I enzymes. Punctual changes in the allosteric binding site of Phe and adjacent region are responsible for the observed relaxed regulation. The phylogeny of plant ADTs evidences that the emergence of type-II isoforms with relaxed regulation occurred at some point in the transition between non-vascular plants and tracheophytes enabling the massive production of Phe-derived compounds, primarily lignin, which are attributes of vascular plants.

## INTRODUCTION

Aromatic amino acids (AAAs) phenylalanine (Phe), tyrosine (Tyr) and tryptophan (Trp) are of paramount importance for all forms of live. However, only bacteria, fungi and plants, have the necessary biochemical pathways for the biosynthesis of these amino acids (Maeda and Dudareva, 2012). In accordance with their importance, evolution has triggered regulatory mechanisms of AAA biosynthesis at the transcriptional and post-transcriptional levels, enabling a fine control of the metabolic flux through these pathways. This is particularly important in plants, where AAAs serve as precursors for the biosynthesis of a wide range of natural compounds including phenylpropanoids, alkaloids, indole auxins and betalains (Schenk and Maeda, 2018). Some of these downstream products, such as lignin, account for a large extent of the total plant biomass (Tohge et al., 2013). An adequate provision of precursors will be necessary to maintain the production of such specialized metabolites.

In plants, the biosynthesis of Phe occurs through two alternative routes (**Figure 1A**). In the arogenate pathway, prephenate is transaminated by prephenate-aminotransferase (PAT) to generate arogenate, which is decarboxylated and dehydrated by arogenate dehydratase (ADT) to give Phe (Bonner and Jensen, 1987). Alternatively, in the phenylpyruvate pathway prephenate is converted first into phenylpyruvate by prephenate dehydratase (PDT), which is the substrate of phenylpyruvate aminotransferase, producing Phe (Yoo et al., 2013). This last pathway has been postulated to be cytosolic in plants (Qian et al., 2019). The arogenate pathway has been reported to be responsible for the main provision of Phe (Maeda et al., 2010; 2011), although various investigations have reported the contribution of the phenylpyruvate pathway (Yoo et al., 2013; Oliva et al., 2017; El-Azaz et al., 2018; Qian et al., 2019). Furthermore, arogenate is a precursor for the biosynthesis of Tyr through the action arogenate dehydrogenase (ADH/TyrAa) (Fischer and Jensen, 1987b; Bonner and Jensen, 1987; reviewed by Schenk and Maeda, 2018).

**Figure 1.**
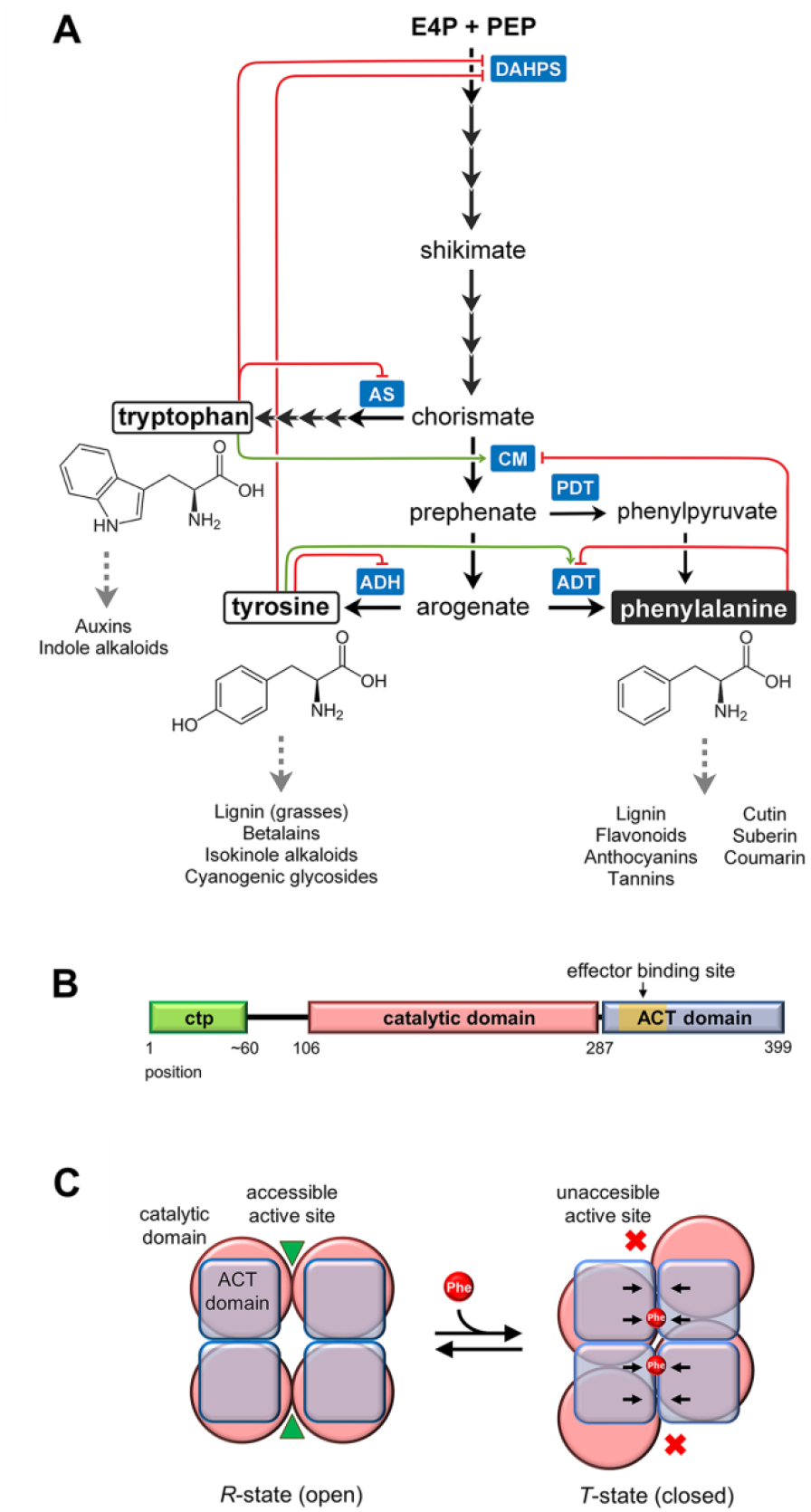
Effector-mediated regulation of the biosynthesis of aromatic amino acids in plants. **A)** Anthranilate synthase (AS), chorismate mutase (CM), arogenate dehydrogenase (ADH) and arogenate/prephenate dehydratase (ADT/PDT) are targets of different feedback-regulatory loops. **B)** Structure of plant ADTs include a N-terminal putative plastid transit peptide (ctp), a catalytic domain and a C-terminal ACT regulatory domain, in which the PAC domain is comprised **C)** Allosteric regulation of ADT by Phe is mediated by a conformational change in the enzyme, postulated as a homotetramer in its quaternary structure. Phe binds to the ACT domain in a pocket formed in the conjunction of the ACT domains from two enzymes monomers. The binding of Phe promotes the reversible transition of the whole enzyme to a T-state, in which the accessibility of the active site is reduced (adapted from Tan et al., 2008).

In addition to transcriptional regulation, some key enzymes of AAA biosynthesis are subjected to effector-mediated regulation mechanisms that determine flux allocation into different branches of the shikimate pathway (Maeda and Dudareva, 2012; **Figure 1A**). ADTs from plants belong to a family of enzymes that are composed of an N-terminal cyclohexadienyl dehydratase catalytic domain fused to an ACT regulatory domain (**Figure 1B**). Cyclohexadienyl dehydratases have the potential of using prephenate and L-arogenate as alternative substrates (Xia et al., 1991; El-Azaz et al., 2016; Clifton et al., 2018), being the superior efficiency in the use of one or other substrate the cause for the enzyme name. These enzymes are typically tetramers (dimers of dimers): dimerization is mediated by the interaction between catalytic and regulatory domains of the monomers, whereas the tetramer is formed only by ACT-ACT contacts (Tan et al., 2008). The ACT domain mediates in the feedback-inhibition of the enzyme by Phe, by inducing a conformational change that makes the active site inaccessible to the substrate (**Figure 1C**) (Jung et al., 1986; Pohnert et al., 1999; Tan et al., 2008). This domain was first characterized in the bacterial enzymes aspartate kinase, chorismate mutase and bifunctional chorismate mutase/prephenate dehydrogenase TyrA, from which its name is derived and is present in a wide range of enzymes that are regulated in response to amino acid levels. Previous investigations have demonstrated that the mutation of the residues involved in the allosteric biding of Phe results in feedback insensitive ADTs, thereby promoting the accumulation of very high levels of Phe in rice and Arabidopsis (Yamada et al., 2008; Huang et al., 2010). The severe effect of ADT-deregulation suggests that effector-mediated regulation of this activity has an important role in controlling Phe homeostasis.

In this study, we report that ADTs from vascular plants are distributed into two groups of isoforms, type-I and type-II, with different levels of feedback-inhibition by Phe. Type-I enzymes, which are common to all land plants and algae ancestors, exhibit a tight inhibition by Phe. Conversely, type-II enzymes show a considerably lesser degree of inhibition by Phe, and are only found in euphyllophytes (ferns and seed plants). Consequently, the overexpression of type-II isoforms resulted in a dramatic accumulation of Phe in the leaves, reaching levels up to 40-times higher than those observed following the expression of type-I enzymes. We have found that the response to Phe as a negative effector is determined by differences in the sequence of the Phe binding site and neighbor regions within the ACT domain. *In vitro* kinetic studies, supported by *in silico* modeling of ADTs from plants and site-directed mutagenesis, suggest that such regulatory differences are due to changes in the affinity towards Phe, along with differences in the inhibition mechanism. Phylogenetic studies of a large number of sequences from lycophytes and ferns support that type-II ADTs diverged from a pre-existent gene duplicate of a type-I isoform in the ancestors of modern vascular plants, probably as an adaptation to the massive demand of lignin and other Phe-derived compounds. Taken together, these findings provide new insights into the biochemical regulation and evolution of Phe biosynthesis in land plants, with possibilities for future biotechnological applications.

## RESULTS

### Two ADTs from maritime pine exhibit radical differences in their sensitivity to Phe inhibition

ADT activity in plants has long been known to be subjected to feedback-inhibition by the end-product of the reaction, Phe (reviewed by Maeda and Dudareva, 2012). Nevertheless, previous works were performed using crude extracts from plant tissues, and provided only a limited information on how this regulatory mechanism affects the various plant ADT isoforms. For this reason, we decided to examine the feedback inhibition of ADT activity by using the recombinant enzymes from maritime pine (*Pinus pinaster*), PpADT-C and PpADT-G, which correspond to the two common clades of ADT sequences from flowering plants (El-Azaz et al., 2016). These two main groups of ADTs present some differences in a 21 amino acids region named PAC (from PDT activity conferring), which overlaps the allosteric Phe binding site and several residues from the tetramerization interface in the ACT domain (**Figure 1B**). The PAC sequence is present in PpADT-G but accumulates some non-conservative changes in PpADT-C, and has been shown to be correlated with the ability of the enzyme to complement PDT deficiency in the yeast auxotrophic mutant *pha2* (El-Azaz et al., 2016).

The kinetic characterization of ADT activity displayed by PpADT-G and PpADT-C showed that both enzymes exhibit a slight positive cooperativity by arogenate with an estimated Hill index (*h*) of 1.4 for PpADT-G and 1.6 for PpADT-C (**Supplemental figure 1**; **Supplemental table 1**). The *S*_0.5_ and *V*_max_ parameters were estimated to be 47.3 μM and 16.0 pKat/μg for PpADT-G and 49.3 μM and 0.3 pKat/ μg for PpADT-C.

PpADT-G reached 50% of inhibition (*IC*_50_) at 27.6 μM of Phe when the initial concentration or arogenate was set at 100 μM (**Figure 2A**; **Table 1**). The inhibition of ADT activity at micromolar levels of Phe was found to be consistent with previous reports (Jung et al., 1986). Surprisingly, no significant decrease in ADT activity was observed for PpADT-C activity up to 100 μM of Phe (Figure 2A), a condition at which PpADT-G was found to be mostly inactive.

**Figure 2.**
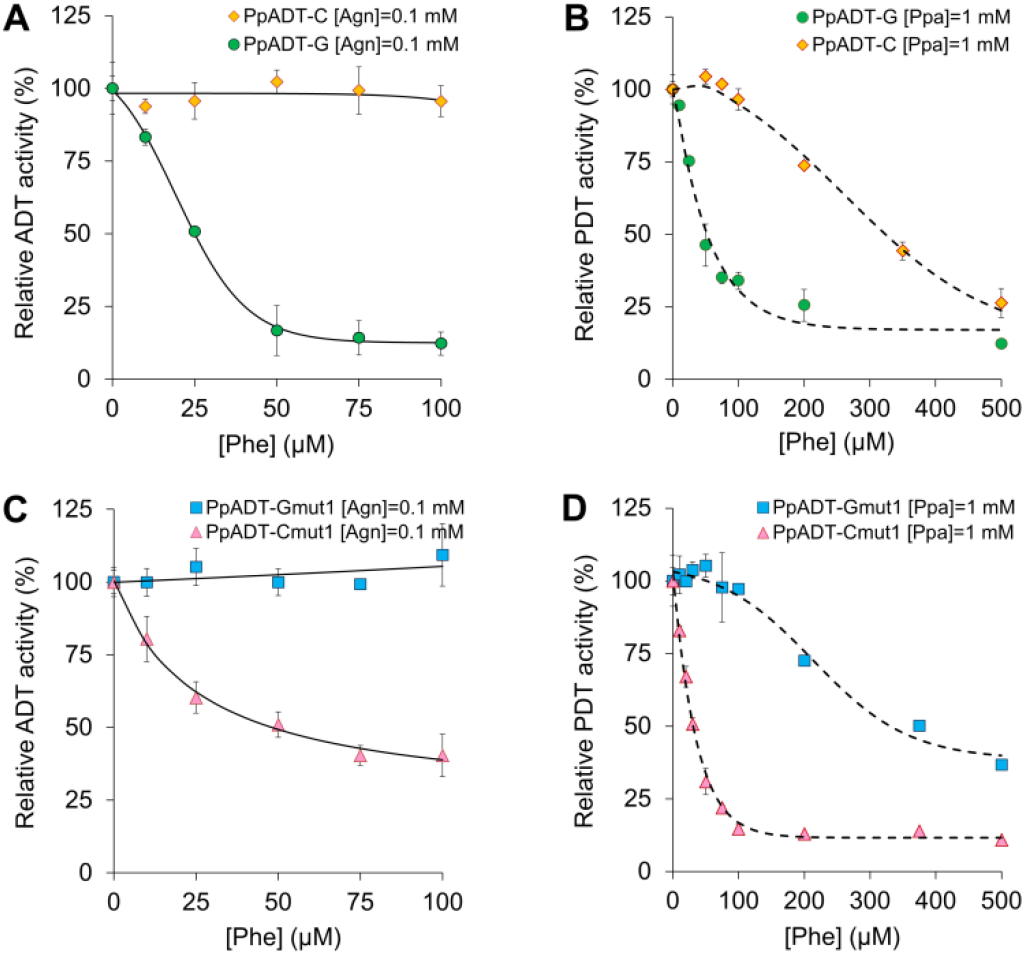
Alternative ADT isoforms have different sensitivities to Phe as negative effector. **A)** Inhibition of ADT activity by Phe in the wild type enzymes PpADT-G and PpADT-C. **B)** Inhibition of PDT activity in PpADT-G and PpADT-C. **C)** Inhibition of ADT activity in the chimeric enzymes PpADT-Gmut^1^ and PpADT-Cmut^1^. **D)** Inhibition of PDT activity in PpADT-Gmut^1^ and PpADT-Cmut^1^. Errors bars represent *SD* from 3 independent replicates. Agn, arogenate; Ppa, prephenate.

**Table 1.** Kinetic parameters of Phe-mediated inhibition in PpADT-G, PpADT-G, PpADT-Gmut^1^ and PpADT-Cmut^1^. Parameters are expressed as an average of three independent replicates ± *SD*. *IC*_50_ values were determined at 0.1 mM of arogenate (for ADT activity) or 1 mM of prephenate (for PDT activity).

|  | ADT | PDT |  |  |
| --- | --- | --- | --- | --- |
|  | *IC*<sub>50</sub> (μM of Phe) | *IC*<sub>50</sub> (μM of Phe) | *K*<sub>i</sub> (μM of Phe) | type of inhibition |
| PpADT-G | 27.6 ± 1.5 | 47.7 ± 3.0 | 28.1 ± 1.0 | uncompetitive |
| PpADT-C | no significant inhibition<br>up to 100 μM of Phe | 320.2 ± 10.7 | n.d. | cooperative |
| PpADT-Gmut<sup>1</sup> | no significant inhibition<br>up to 100 μM of Phe | 389.9 ± 15.4 | n.d. | cooperative |
| PpADT-Cmut<sup>1</sup> | 52.8 ± 2.9 | 36.1 ± 3.4 | 170.5 ± 16.3 | uncompetitive |

To further characterize this differential response to Phe, we decided to take advantage of the PDT activity exhibited by both enzymes, bypassing the technical limitation that the addition of large amounts of Phe represents for accurate determination of ADT activity. Assayed as PDT at 1 mM of prephenate, PpADT-G reached *IC*_50_ at 47.7 μM of Phe (**Figure 2B; Table 1**). Kinetics analysis indicated that the inhibition mechanism of PpADT-G by Phe is apparently uncompetitive, which implies that Phe only can only bind to the enzyme when the enzyme-substrate complex is already established (**Supplemental Figure 2**). The affinity constant for Phe (*K*_i_) was estimated to be 28.1 μM. In contrast, PpADT-C was found to be inhibited only over 100 μM of Phe, with an estimated *IC*_50_ value of 320 μM (**Table 1**). In the absence of Phe, PpADT-C exhibited a Michaelian response to substrate concentration, whereas it progressively switches to sigmoidal kinetics in the presence of increasing amounts of Phe (**Supplemental figure 2**). This behavior is characteristic of allosteric regulation (Palmer, 1995).

### Differences in the Phe binding region determine the relaxed regulation of PpADT-C

PpADT-G and PpACT-C differ in the sequence of the allosteric Phe binding site and oligomerization interface within the ACT domain. To address whether such changes in the sequence explain the different levels of inhibition by Phe, a domain swapping of this sequence motif was performed between PpACT-G and PpACT-C, resulting in two chimeric enzymes: PpADT-Cmut^1^, which contains the PAC domain from PpADT-G, and PpADT-Gmut^1^, its reciprocal counterpart. The response to Phe in the chimeric enzymes was changed compared to their wild-type versions.

PpADT-Cmut^1^ exhibited uncompetitive inhibition (**Supplemental Figure 2**), with estimated *IC*_50_ values of ~52.8 and ~36.1 μM (assayed respectively as ADT or PDT; **Figure 2C and 2D; Table 1**), resembling PpADT-G. The *K*_i_ parameter for Phe was estimated to be 170.5 μM, which was significantly higher than that of PpADT-G.

The reciprocal mutant enzyme PpADT-Gmut^1^ exhibited relaxed regulation in response to Phe (**Figure 2C and 2D; Table 1**). No significant inhibition of ADT activity was observed up to 100 μM of Phe, and the *IC*_50_ value was determined to be almost 400 μM. Moreover, it was also corroborated that the inhibition mechanism changed from uncompetitive to allosteric (**Supplementall Figure 2**).

### Deregulated ADTs are widespread in seed plants

Phylogenetic analysis of the ACT domain of ADTs from seed plants indicates that this domain is distributed into two groups in all vascular plants analyzed (**Figure 3A**). ACT domains that clustered together with the ACT domain from PpADT-G, which contains the PAC sequence, were named type-I. Conversely, those containing the ACT domain from PpADT-C were named type-II, and included enzymes lacking the PAC sequence. Two additional clusters of ADTs, integrated by sequences from conifers and monocots that do not correspond to the features of type-I and type-II isoforms, were also identified. We hypothesized that distribution could correspond to the existence of regulated (type-I) and deregulated (type-II) ADT isoforms among spermatophytes, not only conifers.

**Figure 3.**
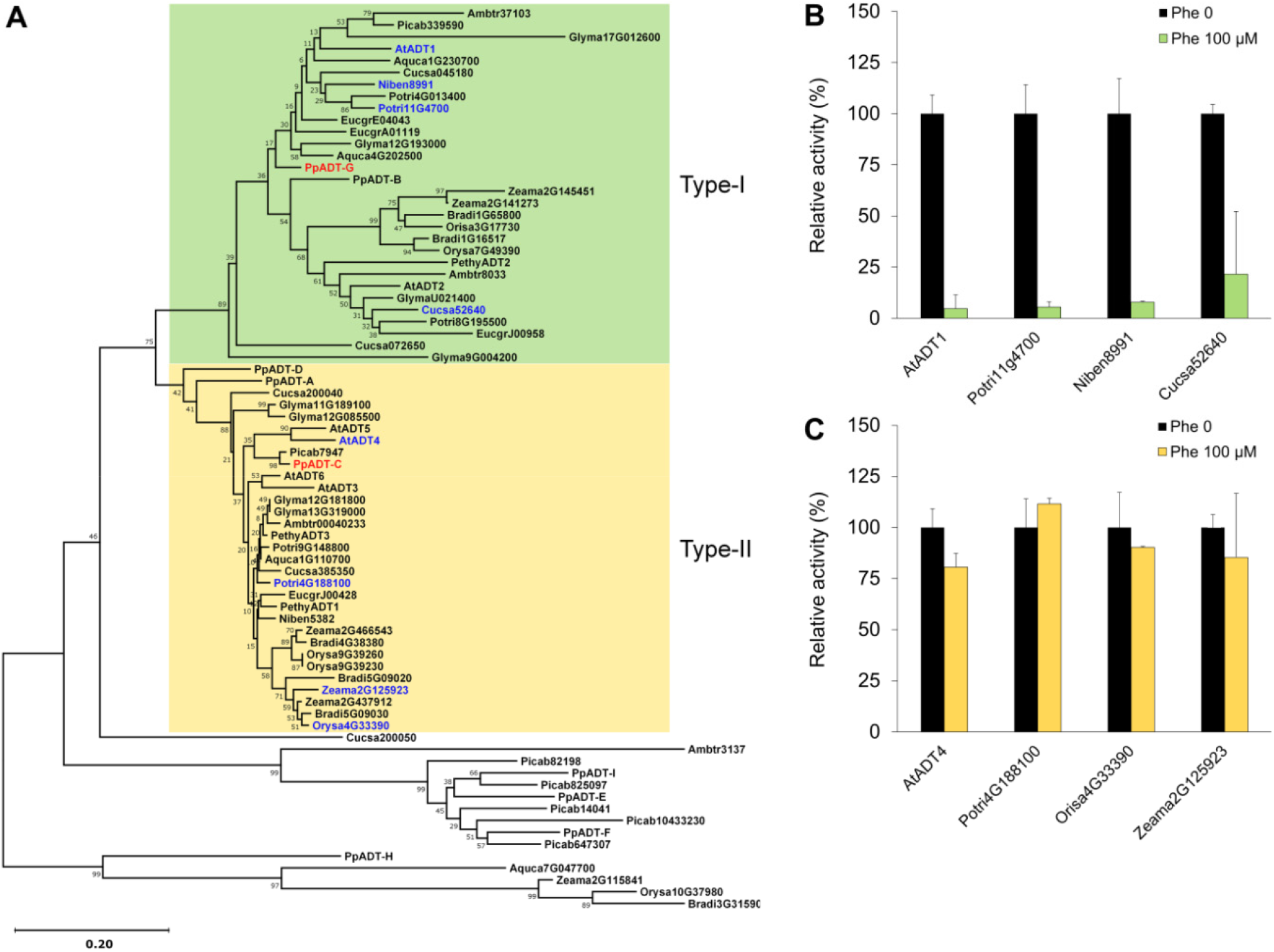
ADTs from seed plants are distributed into two common groups, type-I and II, that differ in the level of regulation by Phe. **A)** Neighbor-joining phylogenetic analysis of the ACT domain of ADTs from seed plants. Type-I, corresponding to putatively tight regulated ADTs, is marked in green. Type-II, corresponding to ADTs with relaxed regulation, is marked in yellow. Tree was set unrooted. Confidence probability is expressed in % and was estimated using the bootstrap test (1000 replicates). Deletion of ambiguous position was set up at a conservation rate of 90%. Species abbreviations: Ambtr, *Amborella trichopoda;* Aquca, *Aquilegia caerulea;* At, *Arabidopsis thaliana;* Bradi, *Brachypodium distachyon;* Cucsa, *Cucumis sativus;* Eucgr, *Eucalyptus grandis;* Glyma, *Glycine max;* Niben, *Nicotiana benthamiana;* Orysa, *Oryza sativa;* Pethy, *Petunia hybrida;* Picab, *Picea abies;* Potri, *Populus trichocarpa;* Pp, *Pinus pinaster;* Zeama, *Zea mays*. **B)** Effect of 100 μM Phe over ADT activity in recombinant type-I ADTs from different plants (names in blue in A). ADT activity is expressed in relative units (control without Phe = 100% activity). **C)** Effect of Phe over type-II recombinant ADTs (names in blue in A). ADT activity was determined in triplicate at 100 μM of substrate (arogenate). Errors bars represent *SD.*

To contrast this hypothesis, we identified and cloned four type-I ADTs from distinct species of flowering plants: *Arabidopsis thaliana* (AtADT1), *Populus trichocarpa* (Potri11G4700), *Nicotiana benthamiana* (Niben8991) and *Cucumis sativus* (Cucsa52640). Such proteins were recombinantly produced, and ADT activity was determined at an initial concentration of Phe (100 μM) and compared to the control (absence of Phe) (**Figure 3B**). The four type-I enzymes exhibited a strong decrease in ADT activity (>90%) at 100 μM of Phe, similar to the type-I enzyme PpADT-G. In parallel, we performed the same experiment by taking four type-II enzymes: AtADT4 (from *A. thaliana*), Potri4G188100 (from *P. trichocarpa*), Orisa4G33390 (from *Oryza sativa*) and Zeama2G125923 (from *Zea mays*). The ADT activity exhibited by the type-II enzymes was not significantly affected by the concentration of Phe employed for the assay (**Figure 3C**), confirming our previous observations in PpADT-C and demonstrating that deregulated ADTs are widespread in flowering plants.

### Overexpression of type-II ADTs has a major impact on Phe levels *in planta*

To test the physiological significance of allosteric deregulation of type-II ADTs, we determined how the overexpression of type-II enzymes would impact Phe accumulation *in planta* compared to type-I enzymes. Leaves from *N. benthamiana* plants that overexpressed the type-I enzymes PpADT-G, AtADT1 and AtADT-2, accumulated Phe to an average level of 4, 32 and 6-times those the control (GFP), respectively. In contrast, the Phe levels in the leaves overexpressing the type-II enzymes PpADT-C and PpADT-A were found to be approximately 145 and 160 times higher compared to the control, and up to 40 times higher compared to the type-I enzymes (**Figure 4**). The levels of the enzymes were determined by western blotting in the same samples (**Supplemental Figure 3**), indicating that the large differences observed in Phe content cannot be attributed to a higher expression level of type-II enzymes.

**Figure 4.**
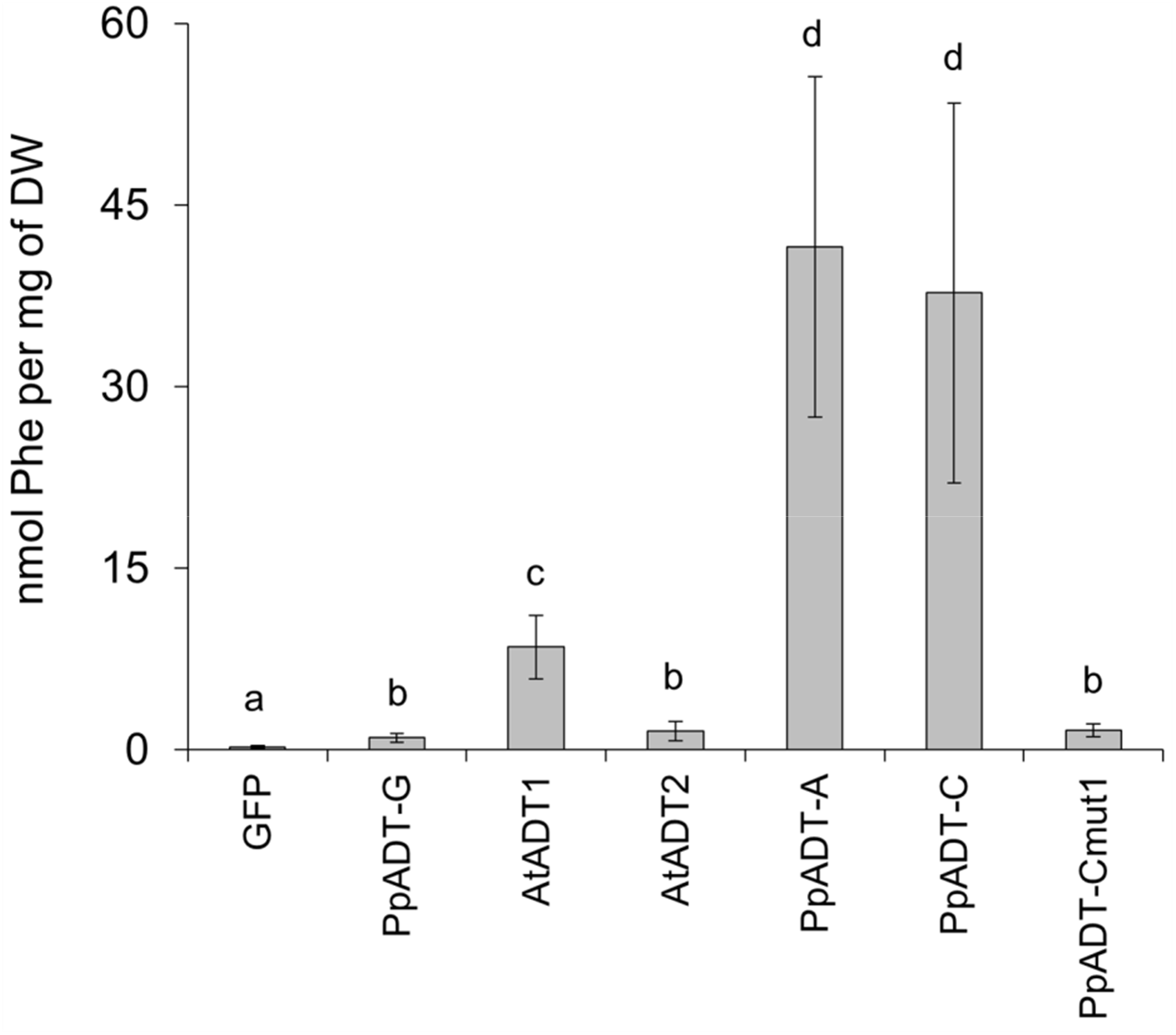
Phenylalanine accumulation in plants overexpressing regulated or deregulated ADTs. Errors bars represent *SD* from 8 independent replicates. Different letters indicate significant differences in the Student’s t-test (*P*-value 0.01). Species abreviations: *Pp*, *Pinus pinaster*; *At*, *Arabidopsis thaliana*. GFP, green fluorescent protein.

To further support the essential role of the PAC domain in the allosteric regulation of ADTs, the chimeric enzymes PpADT-Gmut^1^ and PpADT-Cmut^1^ were included in this experiment. Precedent *in vitro* characterization indicates that the relaxed regulation by Phe can be exchanged by swapping the PAC domain. Consistently, leaves overexpressing PpADT-Cmut^1^ accumulated Phe to a level comparable to leaves overexpressing wild-type PpADT-G (**Figure 4**). The estimation of the protein levels of PpADT-Cmut^1^ by western blot analysis indicated that, in this regard, no major differences could be found when compared to the wild type version (Supplemental Figure 3). After several attempts, the expression of PpADT-Gmut^1^ was found to be highly irregular and undetectable in most of the samples (Supplemental Figure 3). Overall, differences in Phe accumulation are likely a consequence of different sensitivity of the enzymes to Phe as a negative effector.

### Various residues within or adjacent to the Phe binding pocket in the ACT domain determine a lower sensitivity to Phe as a feedback-inhibitor

Phylogenetic analysis differentiates two groups of ADTs in seed plants with a different sequence in the region that is responsible for the allosteric binding of Phe. Type-I isoforms have Thr/Ser^303^, Leu^304^, Pro^308^, Gly^309^, Ala^314^, Ala^316^, Val^317^, Leu^320^ and Asn^324^, with few exceptions (positions as in PpADT-G; **Supplemental Figure 4**). In contrast, Ala^303^, His/Gln^304^, Thr^308^, Ser^309^, Val^314^, Ser^316^, Ala^317^, Phe^320^ and Ser^324^ are common features in type-II enzymes (**Supplemental Figure 4)**. **Figure 5A** depicts a logo sequence of this region that summarizes such differences between type-I and type-II ADTs.

Therefore, we performed a phylogeny-guided site directed mutagenesis study in PpADT-G. Differences in the consensus sequence between type-I and type-II isoforms were used to generate 11 mutant versions of the PpADT-G (**Figure 5B**), and the apparent *IC*_50_ values were determined in their recombinant forms (**Table 2**). *IC*_50_ was found to be significantly increased in the mutant proteins PpADT-Gmut^6^, Gmut^62^, Gmut^8^, Gmut^10^, Gmut^101^ and Gmut^102^, ranging from 184 μM for PpADT-Gmut^10^ to 445 μM in PpADT-Gmut^8^ (**Figure 5C**). On the other hand, mutations affecting the residues Ala^314^, Ala^316^, Val^317^ and their combinations (mutant proteins PpADT-Gmut^9^, Gmut^11^, Gmut^12^, Gmut^13^, and Gmut^16^) increased the sensitivity towards Phe as inhibitor (**Supplemental Figure 5**). These results indicate that different residues between positions 303 and 324 of PpADT-G contribute to modulating the sensitivity to feedback-inhibition by Phe.

**Figure 5.**
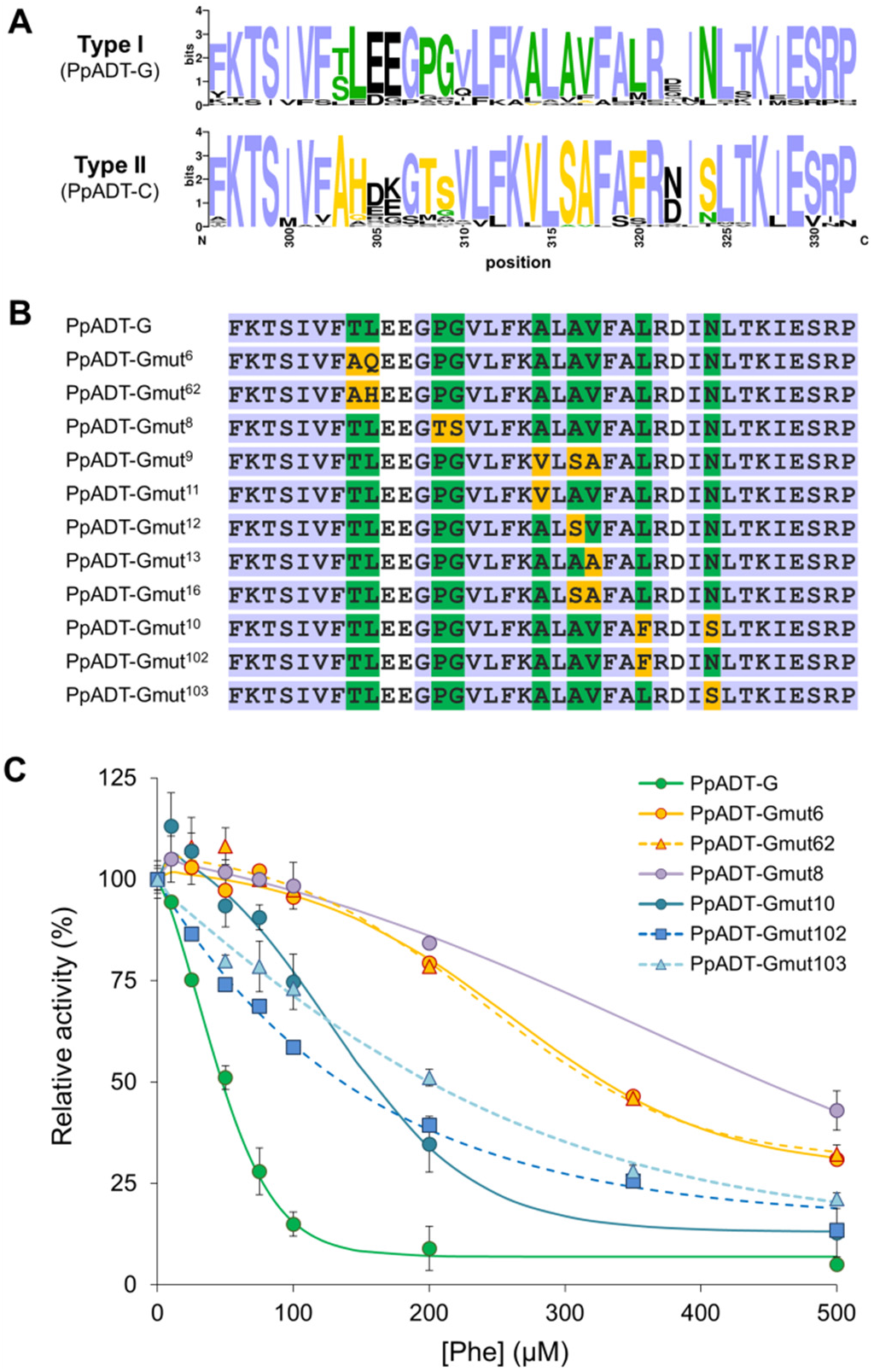
Decreased sensitivity to feedback-inhibition by phenylalanine is determined by various residues in the Phe binding region. **A)** Type-I (green) and type-II (yellow) enzymes differ in the primary sequence of the Phe binding pocket region within the ACT domain. Purple color indicates residues that are preserved between both groups; black color corresponds to barely conserved residues. **B)** Site directed mutagenesis affecting PpADT-G, replacing residues that are highly conserved in type-I isoforms (green) by the corresponding residues from type-II isoforms (yellow). **C)** Determination of the *IC*_50_ parameter in the mutant versions of PpADT-G with decreased sensitivity to Phe as a negative effector, compared to wild type. Measurements were done in triplicate at a concentration of prephenate of 1 mM. Error bars represent *SD*.

**Table 2.** *IC*_50_ values for Phe in the mutant enzymes derived from PpADT-G. *IC*_50_ is expressed as an average of three independent replicates ± *SD* at 1 mM of substrate (prephenate).

| | substitution(s) | $IC_{50}$ (μM) |
| --- | --- | --- |
| PpADT-Gmut<sup>6</sup> | Thr<sup>303</sup> → Ala<sup>303</sup><br>Leu<sup>304</sup> → Gln<sup>304</sup> | 332.8 ± 8.0 |
| PpADT-Gmut<sup>62</sup> | Thr<sup>303</sup> → Ala<sup>303</sup><br>Leu<sup>304</sup> → His<sup>304</sup> | 333.5 ± 3.1 |
| PpADT-Gmut<sup>8</sup> | Pro<sup>308</sup> → Thr<sup>308</sup><br>Gly<sup>309</sup> → Ser<sup>309</sup> | 444.8 ± 42.1 |
| PpADT-Gmut<sup>9</sup> | Ala<sup>314</sup> → Val<sup>314</sup><br>Ala<sup>316</sup> → Ser<sup>316</sup><br>Val<sup>317</sup> → Ala<sup>317</sup> | 17.5 ± 3.7 |
| PpADT-Gmut<sup>11</sup> | Ala<sup>314</sup> → Val<sup>314</sup> | 14.6 ± 2.3 |
| PpADT-Gmut<sup>12</sup> | Ala<sup>316</sup> → Ser<sup>316</sup> | 26.0 ± 4.8 |
| PpADT-Gmut<sup>13</sup> | Val<sup>317</sup> → Ala<sup>317</sup> | 23.6 ± 1.4 |
| PpADT-Gmut<sup>16</sup> | Ala<sup>316</sup> → Ser<sup>316</sup><br>Val<sup>317</sup> → Ala<sup>317</sup> | 17.2 ± 0.2 |
| PpADT-Gmut<sup>10</sup> | Leu<sup>320</sup> → Phe<sup>320</sup><br>Asn<sup>324</sup> → Ser<sup>324</sup> | 184.0 ± 24.7 |
| PpADT-Gmut<sup>102</sup> | Leu<sup>320</sup> → Phe<sup>320</sup> | 135.4 ± 0.7 |
| PpADT-Gmut<sup>103</sup> | Asn<sup>324</sup> → Ser<sup>324</sup> | 234.3 ± 4.9 |

### *In silico* modeling provides structural support for the deregulation of type-II ADTs

To elucidate how the differences in the primary structure of the ACT domain can determine the allosteric response to Phe, PpADT-G and PpADT-C 3D-structures were modeled by homology (**Figure 6**). The PAC region is located at the N-terminal side of the ACT regulatory domain (residues 303 to 324 and 332 to 353 in ADT-G and ADT-C, respectively), encompassing the last three residues of the first β-strand of ACT, the following α-helix and the loops at both ends of the helix (**Figures 6A; Supplemental Figure 6**). In the relaxed form, the residues involved in the binding of Phe form four grooves open to the solvent at each edge of the ACT-dimerization interface (two by dimer). In the tense form, the grooves between ACT domains change to close cavities, and the ACT-interfaces between the dimers and tetramers show a tighter fit (**Figure 6A and 6B**). In addition, the catalytic domains in each dimer are closer than in the full active form, changing the active site conformation and reducing the accessibility of substrate (**Figure 6A**; Tan K. et al., 2008).

**Figure 6.**
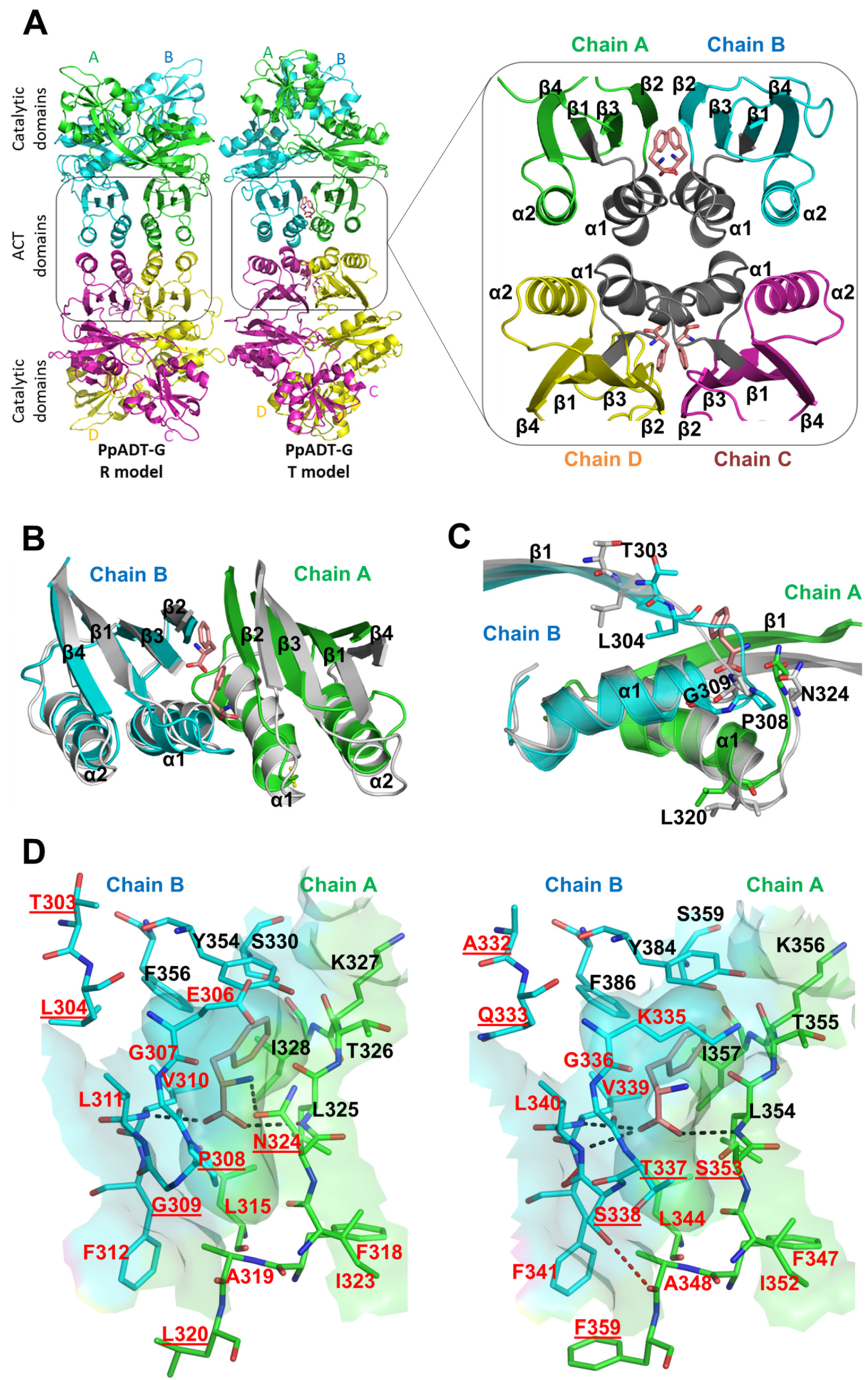
3D-modeling of the allosteric Phe cavity. **A)** Right, cartoon of PpADT-G R and T structural models. Tetramers are colored by chain (A to D), ACT regulatory domains are framed by a rectangle. Allosteric Phe in T model are shown as sticks. A/B and C/D dimers form a tetramer through ACT domain interactions. Rotation of ACT domains, restructuring of the ACT β-sheet in contact with the catalytic domains and active sites closure are remarkable conformational changes between R and T models. Left, detail of the ACT domains of PpADT-G in the tense conformation. The secondary structure elements of the domain are identified (β1α1β2β3α2β4 fold). In grey, the PAC region in each monomer, encompassing the β1 C-terminal end, β1/α1 connecting loop, α1 and half of α1/β2 connecting loop. The four allosteric Phe are shown in stick representation, two by dimer interface. **B)** Superposition between ACT dimers from tense (colored by chain as in panel A) and relaxed (grey) structural models, highlighting conformational differences. PAC regions from chains A and B would have to approach to form the cavities. **C)** Zoom over panel B showing one of the Phe binding sites. The residues that have been mutated in the present work resulting in the increase of Phe *IC*_50_ are represented as sticks in the tense (green and cyan) and relaxed (grey) conformations. **D)** Transparent surface detail of one of the allosteric cavities from PpADT-G (left) and PpADT-C (right) tense structural models, revealing the bound Phe represented as sticks. Residues forming the cavity from chains A and B are shown as green and cyan sticks, respectively. PAC residues from both chains are named with red letters, whereas black letters name cavity forming residues out of the PAC region. PpADT-G residues that have been mutated to the equivalent residues in PpADT-C are underlined in both enzymes. Thr^303^ and Leu^320^ from PpADT-G (Ala^332^ and Phe^359^ in PpADT-C) are outside but close to the allosteric cavity. Discontinuous black lines represent polar interactions between residues of the cavity and the allosteric Phe backbone. Asn^324^Ser mutation in PpADT-G (Ser^353^ in PpACT-C) avoids the interaction with the backbone amino group of Phe and the cavity fails to close properly. Discontinuous red line shows the polar interaction between Ser^338^ and Ala^348^ in PpADT-C, which is not possible between the equivalent positions in PpADT-G. Amino acid one letter code was used for naming the residues. Red and blue colors were used for oxygen and nitrogen atoms, respectively, in the stick representation.

According to our predictions, Ser^353^ in PpADT-C instead of Asn in the PpADT-G equivalent position (Asn^324^) avoids the polar interaction with the Phe amino group and, due to its smaller size, keeps two of the four Phe binding cavities slightly open to the solvent, which would be compatible with a reduction in affinity for Phe as inhibitor (**Figures 6C and D**). In PpADT-Gmut^10^, this substitution was introduced together with Leu^320^Phe (Phe^359^ in PpADT-C). The interactions between large residues, such as Phe^312^ and Leu^320^ (Phe^341^ and Phe^359^ in PpADT-C) are responsible for tetramerization of the monomers (**Figure 6D**). The transition from the relaxed into the tense state involves a decrease in the distance between such residues. As long as Phe is larger than Leu, steric hindrance between the eight Phe rings placed in the ACT tetramer may restrict the concerted movement to reach the tense conformation (see Morph simulation at the Supplemental material). Therefore, both effects, reduction in the affinity for Phe and less efficient transition to the tense form, are in good agreement with the observed Phe *IC*_50_ increase in Gmut^10^.

Mutations placed at the N-terminal side of the PAC region (PpADT-Gmut^6^, Gmut^62^ and Gmut^8^) have a higher impact on sensitivity to Phe than Leu^320^Phe and Asn^324^Ser (PpADT-Gmut^10^), even though any deleterious effect on the cavity shape or the interactions with the Phe backbone could be inferred from the structural models. Superposition of relaxed and tense models for PpADT-G and PpADT-C shows that residues in this region (303 to 309 and 332 to 338 in PpADT-G and −C, respectively) form a loop connecting the β-strand and the α-helix of the PAC region with a turn, experiencing strong positional and conformational changes in the transition to the tense state (**Figure 6C**). This finding suggests that the flexibility in this loop is important to fulfill this function in the ACT domains. Hence, as long as Gly is the amino acid with the smallest side chain, Gly^309^Ser substitution in PpADT-Gmut^8^ will reduce the flexibility of this loop. Interestingly, a recent study combining *in silico* dynamics simulation and experimental results has shown that this region is part of a ligand gate whose movements allow the entrance of the allosteric Phe to the cavity in the ACT domain of human phenylalanine hydroxylase (Ge et al., 2018). Moreover, it has been found that the same mutation at the homologous position of human phenylalanine hydroxylase (Gly^46^Ser) completely prevents the binding of Phe to the regulatory domain (Leandro et al., 2017). In contrast, Pro^308^Thr (the second mutation in PpADT-Gmut^8^) releases the rigidity in the peptidic backbone, and together with Gly^307^, it likely contributes to smoothing the effect of Gly^309^Ser substitution. In contrast, Thr^308^ (Thr^337^ in PpADT-C) is predicted to form a hydrogen bond with Ala^319^ (Ala^348^ in PpADT-C) from the opposite monomer that forms the cavity (**Figure 6D**), stabilizing the conformation in the tense form. Taken together, our model indicates that Pro^308^Thr and Gly^309^Ser would counterbalance their individual effects to produce a loop that is less flexible but still functional.

Last, PpADT-Gmut^6^ and Gmut^62^ mutations affect Thr^303^ and Leu^304^ at the C-terminal end of the ACT first β-strand (**Figure 6C**). Thr^303^ does not form part of the predicted Phe cavity, whereas Leu^304^ is involved in a small portion of it. In the transition between relaxed and tense conformations, ACT β1 moves together with the loop comprised between positions 303 to 309 to form the cavity (**Figure 6C**; Morph simulation at Supplemental material), changing its backbone interactions with the adjacent strands of the β-sheet and the side chains. Although the Thr^303^Ala mutation does not appears have an effect on the structure or the allosteric conformational change, Leu^304^Gln or Leu^304^His introduce a voluminous side chain with a polar group between hydrophobic residues (**Figure 6D**), producing steric hindrance in the tense conformation, but not in the relaxed conformation. Therefore, the active relaxed conformation would be favored over the tense conformation, in concordance with the increase of *IC*_50_ of PpADT-Gmut^6^ and PpADT-Gmut^62^.

### Deregulated ADTs are a novel and distinctive feature of vascular plants

With the aim of addressing the distribution of deregulated ADT activity through the plant kingdom, we examined the occurrence of the key mutations Thr^303^Ala, Leu^304^His/Gln, Pro^308^Thr, Gly^309^Ser, Leu^320^Phe and Asn^324^Ser, in the PAC domain of ADTs from green algae to flowering plants. We identified 72 non-redundant ADT sequences from green algae, 62 from liverworts, 87 from bryophytes, 78 from lycophytes, 425 from pteridophytes and 69 from spermatophytes. Partial sequences included at least the complete studied region. **Figure 7** summarizes the relative distribution of residues in the Phe binding site and adjacent residues, showing that approximately half of the ADT isoforms from seed plants likely correspond to enzymes recalcitrant to feedback inhibition. Remarkably, the residues that were demonstrated to determine a reduction in the inhibition by Phe are completely absent or highly uncommon in the enzymes from algae and, especially, non-vascular plants (liverworts and bryophytes). This rule can be extended to the enzymes from lycophytes, the most primitives from the extant vascular plants, with the exception of the Thr/Ser^303^→Ala^303^ mutation. In contrast, ADT isoforms from pteridophytes already exhibit the entire set of key mutations that characterize the Phe binding region of type-II isoforms, except for Asn^324^Ser, which is rare. Additionally, the different mutations were found to co-occur within the same isoforms from ferns, similar to seed plants (**Supplemental Figure 5**).

**Figure 7.**
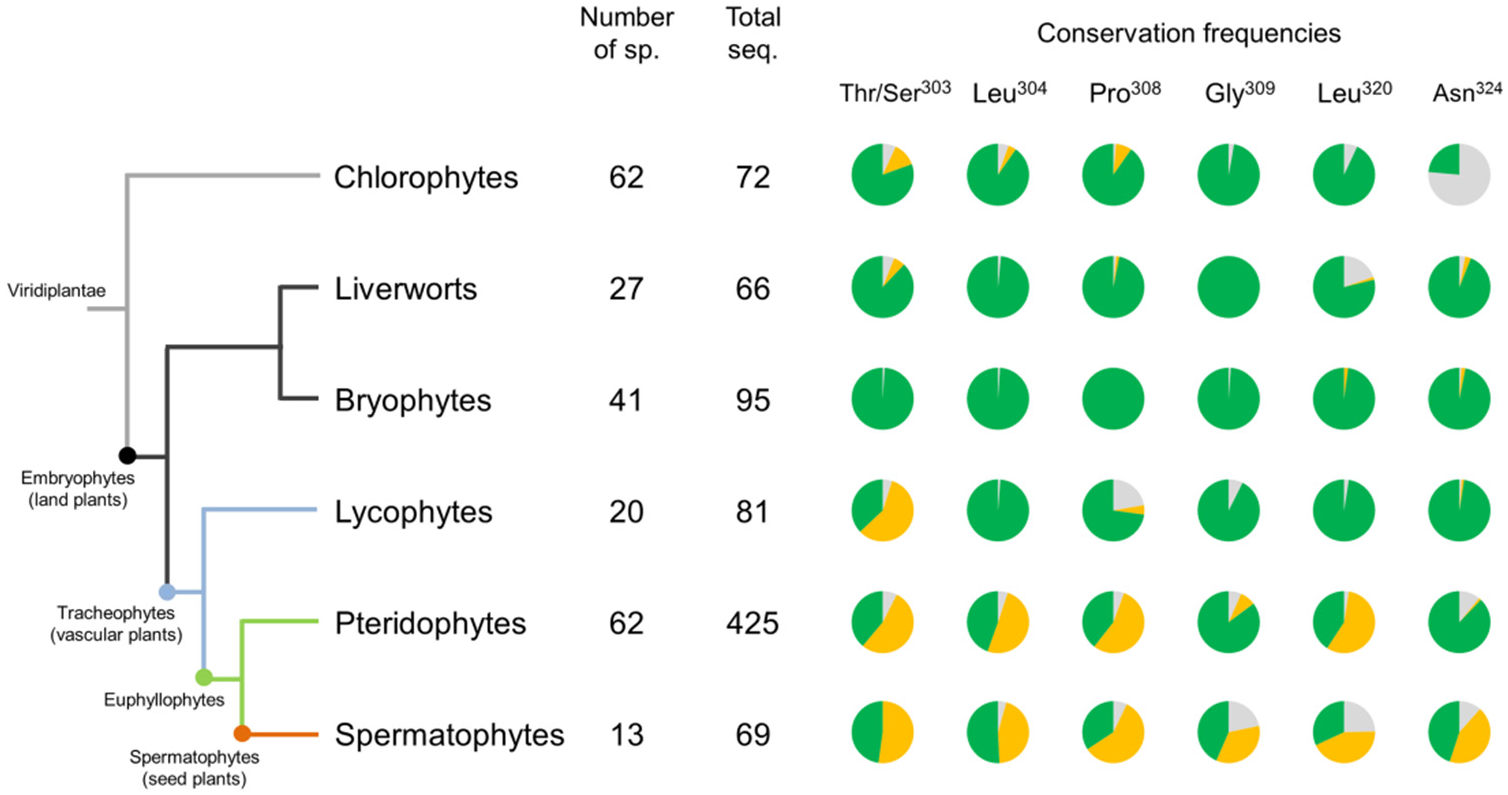
Conservation frequencies in % of critical residues determining high sensitivity towards Phe inhibition of ADT activity. Sequences of ADTs from algae to flowering plants were analyzed. The number of species (Number of sp.) and the number of ADT sequences identified (Total seq.) were indicated for each taxon. The conservation of the residues listed, or a chemically similar residue, that characterize type-I ADTs and determine high sensitivity towards Phe as negative effector, were indicated in green. Yellow color indicates the substitution ratio of these residues by their equivalent counterparts in type-II ADTs or similar residues, promoting a relaxed regulation by Phe. Grey color indicates the occurrence of alternative residues.

The phylogenetic analysis of the ADTs from the two fern species fully sequenced up to date, *Azolla filiculoides* and *Salvinia cucullata* (Li et al., 2018), confirmed that priteridophytes hold type-II isoforms (**Supplemental Figure 7**). Conversely, the ADTs from the three non-vascular plants with available genomes, *Marchantia polymorpha*, *Physcomytrella patens* and *Sphagnum phallax*, were found to cluster outside the type-II group. Very interestingly, sequences from the lycophyte *Selaginella moellendorfii* were found to occupy an intermediate position between type-I and Type-II enzymes. A more detailed examination of the two lineages of ADT isoforms revealed that genes encoding for type-I enzymes have retained an intron-exon structure from the green algae ancestors, whereas genes encoding for type-II isoforms typically lack introns, including type-II enzymes from pteridophytes (**Supplemental Figure 8**). Intronless ADT coding genes are also present in *M. polymorpha*, *P. patens*, *S. phallax* and *S. moellendorfii*, despite the lack of typical type-II enzymes. Overall, these results suggest that deregulated ADTs emerged in the lineage of the Euphyllophytes from a precedent gene duplication event affecting all the extant land plants, which was accompanied by an intron-losing phenomenon.

## DISCUSSION

Effector-mediated regulation of enzymatic activity is a key control mechanism to maintain amino acid homeostasis (**Figure 1**). In the present work, we report that type-I and type-II ADT isoforms from spermatophytes differ in their response to Phe as a negative effector. Type-I isoforms, which are more closely related to enzymes from algae and bacteria, exhibit tight inhibition by Phe, the product of the reaction. Previous literature reported *IC*_50_ values for ADT activity at approximately 35 μM of Phe in crude plant extracts (Jung et al., 1986). These reports are similar to our observations for the recombinant type-I enzyme PpADT-G (**Table 1; Figure 2**). In contrast, type-II ADTs remain active at relatively high Phe levels, as shown for recombinant PpADT-C and other enzymes from various plants (**Figures 2 and 3**). Consequently, the overexpression of type-II enzymes in plant leaves promotes the accumulation of considerably higher levels of Phe than the overproduction of type-I-enzymes (**Figure 4**). Interestingly, the overexpression of AtADT4, a type-II enzyme, was previously demonstrated to have a strong impact on anthocyanin accumulation, as a consequence of reduced sensitivity to Phe inhibition (Chen et al., 2016). Phylogeny- and structure-guided mutagenesis studies have identified a set of residues, from the Phe binding region in the ACT regulatory domain of the enzyme, that are involved in the decreased inhibition observed in type-II enzymes. Phylogenetic evidence indicates that type-II ADTs emerged from type-I isoforms at some point in the evolution of tracheophytes, with a foreseeable impact on the massive production of Phe-derived compounds that takes place in this group of plants.

Our description of a novel clade of ADT enzymes with relaxed feedback inhibition is in keeping with previous reports affecting other key enzymes of AAA biosynthesis in plants. For example, anthranilate synthase (AS) has two isoforms in flowering plants, named as constitutive and inducible, that differ in their sensitivity towards Trp as inhibitor. Constitutive AS, which is expressed at basal levels in different plant tissues, has a *K*_i_ for Trp in the range from 2 to 3 μM. On the other hand, the inducible isoform of AS, which is expressed in response to certain stimuli that presumably involve the synthesis of major amounts of Trp, is notably less sensitive to Trp as negative effector (*K*_i_ from 100 μM to 300 μM; Bohlmann et al., 1996; Song et al., 1998). CM, the commitment step enzyme that channels chorismate into the biosynthesis of Phe and Tyr, is inhibited by both amino acids, whereas Trp promotes its activation (reviewed by Maeda and Dudareva, 2012). The inhibition of plastidial CM by Phe seems to be highly divergent between different groups of plants: *K*_i_ has been estimated to be 1.1 mM in *Papaver somniferum* (Benesova and Bode, 1992) and 550 μM in *Solanum tuberosum* (Kuroki and Conn, 1988), whereas the reported *IC*_50_ values were 50 μM in *Arabidopsis thaliana* (Westfall et al., 2014), 82 μM in *Amborella trichopoda* (Kroll et al., 2017), and 2.6 and 7.4 mM for the two isoforms found in *Physcomitrella patens* (Kroll et. al., 2017). Nevertheless, flowering plants possess a cytosolic isoform of CM, which is insensitive to feedback regulation by AAAs (Eberhard et al., 1996; Westfall et al., 2014). Relative to Tyr biosynthesis, it has been shown that plants in the order Caryophyllales have developed an ADH isoform with relaxed feedback inhibition by Tyr, in a close evolutionary relationship with the production of Tyr-derived betalain pigments in many species from this order (López-Nieves et al., 2018). The occurrence of these deregulated enzymes highlights that in many cases the production of specialized metabolites in higher plants has evolved to provide a surplus of precursors from the primary metabolism.

The kinetic characterization of PpADT-G and PpADT-C indicates the existence of major differences in the mechanism underlying feedback inhibition by Phe. In PpADT-G, we observed that Phe produces a decrease in both *K*_m_ and *V*_max_, an effect that is characteristic of uncompetitive inhibition. The inhibitor is only able to bind the enzyme-substrate complex but not the free enzyme. In PpACT-C, the enzyme exhibits a sigmoidal response to substrate concentration in the presence of Phe. The inhibitor produces a decrease in the apparent affinity of the enzyme for the substrate, with the rate of reaction decreasing at low substrate concentrations This behavior resembles competitive inhibition, although this term is not strictly applicable in this case, as in the presence of exogenous Phe the enzyme does not obey Michaelis-Menten equation (**Supplemental Figure 2**). Previous works suggested that Phe-mediated inhibition of ADT activity in spinach chloroplast extracts was most likely competitive or mixed, which is in close agreement with our findings (Jung et al., 1986). Differences in the mechanism of feedback-inhibition of both types of ADTs could be of major physiological relevance. Phe-induced cooperativity of PpADT-C would allow the recovery of high-speed rates when substrate accumulates, even in the presence of the inhibitor, which would not happen with uncompetitive inhibition of PpADT-G. This result could have a strong impact over Phe levels *in planta*, as observed in **Figure 4**. The *in silico* modeling of the enzymes (**Figure 6**) predicted that deregulation of type-II ADTs is of multifactorial origin: a decreased affinity towards Phe, changes in the flexibility of the peptidic backbone in the PAC region that would impact dynamics of the transition between the relaxed and the tense forms, and a different conformation and stability of the allosteric cavity in the tense form due to steric hindrance. Factors affecting the architecture and dynamics of the allosteric cavity could underlie the different mechanism of inhibition observed between PpADT-G and PpADT-C. On the other hand, the interchange of the PAC region between PpADT-G and PpADT-C improves not only PpADT-C sensitivity to Phe, but also the apparent affinity for prephenate (El-Azaz et al 2016). It suggest, along with enzyme’s folding prediction, that PpADT-C has less active site plasticity and is in general a more rigid molecule than PpADT-G. Type-I PAC sequence could confer a more flexible and dynamic R conformation, increasing active site plasticity for the use alternative substrates (arogenate or prephenate). Additionally, Leu^320^Phe mutation, that increases Phe *IC*_50_, provides first evidence that the tetramer, and not only the dimer, could be important to achieve the tense conformation of the enzyme (Tan et al 2008).

An in-depth evolutionary study of the ADT family indicates that deregulated ADTs from type-II are widespread in seed plants, but absent in green algae, bryophytes and liverworts (**Figure 7; Supplemental Figure 5**). Our study indicates that pteridophytes, the sister group of seed plants (Li et al., 2018), have unequivocally type-II ADTs. The analysis of 425 sequences from ferns revealed that the key substitutions Thr^303^Ala, Leu^304^His/Gln, Pro^308^Thr, and Leu^320^Phe, which contribute to increasing the *IC*_50_ of the enzyme, have similar frequencies to seed plants (**Figure 7**). These mutations usually co-occur in the same enzyme, typically encoded by genes without introns in both pteridophytes and spermatophytes. Lycophytes, the most primitive of the extant vascular plants, are a key group to track the emergence of type-II isoforms. In these plants, isoforms with Thr^303^Ala were observed frequently, as well as Ala^314^Val, Ala^316^Ser, Val^317^Ala, being encoded by intron-less genes (**Supplemental Figure 8**). These last residues, although being a characteristic of type-II ADTs, were shown to decrease the *IC*_50_ of the model enzyme PpADT-G (**Supplemental Figure 6**). The remaining mutations that define type-II enzymes, especially those predicted to have important structural impact according to our model (**Figure 6**), were found to be extremely uncommon in lycophytes. It is unlikely that ADTs from this group are de-regulated isoforms. Based on the evidence provided, we propose that the primitive condition of ADTs in the Viridiplantae lineage was a high sensitivity to feedback inhibition by Phe, as observed in type-I isoforms. The structure of the *ADT* gene family across land plants indicates that this family suffered an early duplication event that affected all embryophytes, accompanied by the loss of introns. These duplicates were retained in vascular plants, and diverged into type-II ADTs at some point after the separation of modern lycophytes and pteridophytes, probably in a stepwise accumulation of key mutations affecting regulatory properties.

The reduction of the allosteric control of Phe biosynthesis has obvious consequences for vascular plants. As metabolism of phenylpropanoids, and particularly lignin biosynthesis, emerged and diversified, the demand of Phe for supplying such downstream pathways dramatically increased. A feasible hypothesis would be that type-I ADTs were not able to properly respond to the increasing demand for Phe in the incipient vascular plants, as far as these enzymes are inhibited when Phe accumulates at relatively low levels. This limitation seems particularly striking when we consider that the bulk of Phe biosynthesis takes place within a confined subcellular compartment, the plastids (Jung et al., 1986; Rippert et al., 2009). As the biosynthesis of phenylpropanoids occurs in the cytosol, Phe must be exported towards this compartment to be further metabolized, a process that has been found to be a major limiting factor in lignin biosynthesis rates (Guo et al., 2018). Hence, an ADT efficiently inhibited at low levels of Phe would remain mostly inactive in the limited space of plastid stroma, where Phe cannot be readily transformed into downstream products, and exportation across the plastid membrane is limiting. Along the evolution of land plants, the inconvenience of a tight feedback inhibition of ADT activity would have favored the stepwise accumulation of mutations in previously duplicated ADT genes, reducing sensitivity to Phe as negative effector. Partially deregulated ADT isoforms would overpass the restrictive allosteric control of the pathway, making them more amenable for sustaining high Phe biosynthesis rates able to fueling a range of new evolved pathways. This hypothesis is supported by previous studies in diverse plant species, indicating that type-II enzymes have a highlighted role in the biosynthesis of Phe-derived compounds. However, the disruption of AtADT4 and AtADT5 in *A. thaliana* has the largest impact on reducing lignin accumulation (Corea et al., 2012). Additionally, in *Arabidopsis*, the overexpression of AtADT4 resulted in Phe hyperaccumulation and elevated levels of anthocyanins, an effect that was not observed when type-I enzymes were overexpressed (Chen et al., 2016). Likewise, *Petunia* x *hybrida* ADT1 has been identified as the major contributor to Phe-derived volatile emission in the petals (Maeda et al., 2010). Finally, in *P. pinaster*, we found that the expression of *PpADT-A* is strongly induced in response to the transcriptional reprograming that leads to the formation of compression wood, a specialized vascular tissue enriched in lignin (El-Azaz et al., submitted).

In conclusion, our results indicate that vascular plants progressively developed a new clade of ADT isoforms with relaxed feedback inhibition. Reduced regulation of the ADT activity must have a huge impact on the biosynthesis of lignin and other phenylpropanoids. Moreover, the identification of sequence motifs responsible for this trait provides an interesting biotechnological target that could help to rationally engineer the production of AAAs and their derived compounds.

## MATERIALS AND METHODS

### DNA constructs

The cloning of the coding region from PpADT-A, PpADT-C, PpADT-G, AtADT1, and AtADT2 was described previously (El-Azaz et al., 2016; 2018). Additional ADTs from *P. trichocarpa* (Potri11G4700, Potri4G188100), *N. benthamiana* (Niben8991). *A. thaliana* (AtADT4), *C. sativus* (Cucsa52640), *O. sativa* (Orisa4G33390) and *Z. mays* (Zeama2G125923) were cloned from cDNA of the corresponding plants (see **Supplemental Table 2** for primer list). All constructs for protein heterologous expression in *E. coli* were cloned into pET30b using the NdeI / NotI sites. Putative plastid transit peptide was removed and C-terminal poli-His tag was added. Mutant chimeric proteins PpADT-Gmut^1^ and PpADT-Cmut^1^ were generated by fusion PCR as described previously (El-Azaz et al., 2016). PpADT-Gmut^62^, Gmut^102^ and Gmut^103^ were generated by site directed mutagenesis using the construct pET30b-PpADT-G as mold (see **Supplemental Table 2** for primers). The generation of the remaining mutant versions of PpADT-G by site directed mutagenesis was described in El-Azaz et al., 2016. Plant expression constructs were cloned into the Gateway^®^ vector pDONR™207 and recombined into pGWB11 (CaMV P35S promoter, c-terminal FLAG^®^ tag; courtesy of Dr. Tsuyoshi Nakagawa, Department of Molecular and Functional Genomics, Shimane University, Japan).

### Protein production and purification

Recombinant ADTs were expressed in *E. coli* strain BL21 DE3 RIL. After optic density at 600 nm reached 0.5-0.6, cultures were chilled on ice for 10 minutes before adding IPTG to a final concentration of 0,5 mM. Induced cultures were incubated for 18 to 20 hours at 12 °C under gentle shaking (~75 rpm). Pellets were collected by centrifugation and preserved frozen at −20 °C. Poly-His tagged recombinant proteins were purified using a nickel resin (Protino Ni-TED 2000 Packed Columns, Macherey-Nagel) and exchanged to Tris buffer 50 mM pH 8.0 in Sephadex^™^ G-25 M resin (PD-10 Columns, GE Healthcare).

### Enzyme assays

PDT activity was determined based on the method described by Fischer and Jensen (1987a). Enzyme assays were performed in Tris buffer 50 mM pH 8.0 with around 5 μg of purified recombinant protein in a final reaction volume of 50 μL. After incubation, the reactions were stopped with the addition of 950 μL of NaOH 2M, mixed immediately, and left to settle for at least 10 minutes at room temperature. Absorbance was registered at 321 nm. Authentic phenylpyruvate was used to do a calibration curve at this wavelength. Prior to enzyme assays, prephenate content in the commercial preparation was estimated by treatment with HCl 1N during 15 minutes at room temperature, to produce its spontaneous decarboxylation to phenylpyruvate, followed by the quantification of this last compound as described before.

ADT activity was determined in a reaction mixture consisting of ~1 μg of purified recombinant protein in Tris buffer 50 mM pH 8.0, to a final volume of 50 μL. Arogenate was prepared as described by El-Azaz et al. (2018), based on the previous protocol published by Rippert and Matringe (2002). Reactions were stopped with 20 μL of methanol, centrifuged at top speed for 5 minutes and filtered through a 0.22 μm nylon filter. Phe production was determined by UPLC/MS using a Waters^®^ Acquitiy UPLC system coupled to a triple quadrople detector (Waters^®^). 1 μL of the sample was subjected to separation using an Acquity UPLC BEH C18 column (1.7μm; 2.1×50 mm) at a flow rate of 0,3 mL/min at 5 °C, in the following gradient: 1 minute in 0.12% acetic acid in water, 3 minutes in a linear gradient from the later solution to methanol 100%, and 1 minute in methanol 100%. Column was rinsed for 2 minutes with 0.12% acetic acid in water between samples. The column effluent was analyzed by positive electrospray ionization under the following settings: capillary voltage 2,45 KV, cone voltage 10 V, source temperature 150 °C, and desolvation temperature 400 °C. Identification and quantification of Phe was based in a calibration curve of pure Phe.

All reactions were incubated at 30 °C in gentle agitation. Enzymes were pre-incubated for 5 minutes before starting the reaction with the addition of the substrate (prephenate or arogenate). When the assay was performed in the presence of Phe as negative effector, this amino acid was added to the initial reaction mixture and pre-incubated with the enzyme for 5 minutes. In all cases, measurements were done at least in duplicate, and the linearity of the reaction was corroborated for a minimum of 5 minutes after starting the reaction. All kinetic adjustments and parameters were obtained by using the Solver tool included in Microsoft Excel.

### Overexpression of ADTs *in planta*

Full-length coding regions were cloned into the Gateway^®^ destination vector pGWB11 (courtesy of Dr. Tsuyoshi Nakagawa, Department of Molecular and Functional Genomics, Shimane University, Japan) following standard procedures. Overexpression constructs were transformed into *Agrobacterium tumefaciens* strain C58C1. Saturated cultures were adjusted to an optic density at 600 nm of 0.25, and were combined with an equal density of a culture carrying the p19 construct (Voinnet et al., 2003). Each 5-weeks-old *N. benthamiana* plant was infiltrated, approximately between 2 to 4 hours before the end of the light period, with four different bacterial clones distributed into the halves of two fully-expanded leaves. Infiltration pattern was rationally designed to equally distribute 16 replicates for each overexpression construct between the plants included in the experiment. Samples were collected 72h after infiltration, and frozen immediately in liquid nitrogen. Leave major nerves were excluded from the sampling to avoid distortion of the dry weight.

### Metabolite extraction

Plant samples were grinded in liquid nitrogen and combined in 8 pairs of replicates for each transgenic protein. Around 100 mg of frozen powder was lyophilized at −40° C during 48 hours. De-hydrated powder was resuspended in 400 μM of extraction buffer (2-amino-2-methyl-1-propanol 0,5% pH 10.0 in EtOH 75%; Qian et al., 2019) and incubated overnight in the cold under vigorous shaking. After incubation, tubes were centrifuged for 5 minutes at 20 000 *g* and 300 μL were recuperated from the supernatant, vacuum dried, dissolved in 100 μL of purified water and frozen at −80° C until analysis. High-grade commercial Phe (20 nmol) was added as internal standard for estimating recovery rate in the control samples, which was around 75%. Phe levels were determined by UPLC/MS as described in the previous section for ADT activity. Phe content was multiplied by 4/3 to correct the estimated recovery rate, and normalized to the dry weight of the sample.

### Protein extraction and western blot analysis

Total proteins from plants were extracted in buffer A (Tris buffer 50 mM pH 8.0, glycerol 10%, SDS 1%, EDTA 2 mM and β-mercaptoethanol 0,1%). Around 100 mg of frozen powder was resuspended in 200 μL of buffer A at room temperature. Samples were centrifuged at 20 000 *g* for 5 minutes and 80 μL was recovered from the supernatant, mixed with 20 μL of 5X Laemmli buffer and denatured at 100 °C for 5 minutes. The remaining supernatant was subjected to removal of excess SDS following the method described by Zaman and collaborators (Zaman et al., 1979), prior to determination of the protein concentration using a commercial Bradford reagent (Bio-Rad^®^ protein assay).

For immunodetection of transiently expressed ADTs in the protein extracts, 30 μg of total proteins was separated by SDS-PAGE electrophoresis. Western blot analysis was used following standard procedures. Transgenic proteins were detected taking advantage of the FLAG^®^ tag included in the construct, using an specific commercial antibody (OctA-Probe mouse monoclonal antibody, Santa Cruz Biotechnology) at 1:500 dilution.

### Homology molecular modeling and model analysis

PpADT-G and PpADT-C three-dimensional models with and without L-Phe bound to ACT regulatory domain were obtained by homology modelling using the software package Modeller 9v11 (Eswar et al., 2006; http://salilab.org/modeller). X-ray crystallographic structures from *Staphylococcus aureus* and *Chlorobium tepidum* PDTs (PDB codes 2qmw and 2qmx) were used as template for the active and L-Phe inhibited states of the enzyme, respectively (Tan et al., 2008). Protein sequence alignment of PpADT-G and PpADT-C with each template used for the homology modelling (see Supplemental material) was constructed on the basis of the alignment of all the available *P. pinaster* ADTs isoforms (from −A to −I; El-Azaz et al., 2016) and both bacterial templates using Clustal omega (Sievers and Higgins, 2014). Ten models were generated for each protein. DOPE (Discrete Optimized Protein Energy) score was used to select the bests models (Shen and Sali, 2006), which were submitted to Molprobity (Chen et al., 2010; http://molprobity.biochem.duke.edu/) for additional verification of their stereochemical quality. PyMOL was used for visualization of the structures, mutation simulation and imaging generation. Morph simulations of the conformational changes between active (R) and inhibited (T) forms upon L-Phe binding were realized using Chimera (Pettersen et al., 2004). R and T 3D-models for PpADT-G and C and Morph simulations for the conformational changes are available as downloadable files in supplementary material in pdb format.

## ACKNOWLEDGEMENTS

We are grateful to Dr. Tomás Vigal García and Virginia Medina Álvarez, from the Laboratorio de Técnicas Instrumentales, Universidad de León (León, Spain) for setting up and performing Phe quantification by UPLC/MS, along with their helpful advice on experimental design and samples preparation for such analysis. This work was supported by grants from the Spanish MICINN (BIO2015-69285-R and RTI2018-094041-B-I00) and Junta Andalucía (Research Group BIO-114).

## AUTHOR CONTRIBUTIONS

JEA: designed the research, performed research, wrote the paper with contributions of all authors; FMC: designed the research, project leader; BB: performed computational analysis; CA: designed the research, project leader; FT: designed the research, performed research.

